# A chemical screen identifies a link between lipid metabolism and mRNA translation

**DOI:** 10.1101/2020.05.11.088005

**Authors:** Alba Corman, Dimitris C. Kanellis, Maria Häggblad, Vanesa Lafarga, Jiri Bartek, Jordi Carreras-Puigvert, Oscar Fernandez-Capetillo

## Abstract

mRNA translation is one of the most energy-demanding processes for living cells, alterations of which have been frequently documented in human disease. Using recently developed technologies that enable image-based quantitation of overall translation levels, we here conducted a chemical screen to evaluate how medically approved drugs, as well as drugs that are currently under development, influence overall translation levels. Consistent with current knowledge, inhibitors of the mTOR signaling pathway were the most represented class among translation suppresors. In addition, we identified that inhibitors of sphingosine kinases (SPHKs) also reduce mRNA translation levels independently of mTOR. Mechanistically this is explained by an effect of the compounds on the membranes of the endoplasmic reticulum, which activates the integrated stress response (ISR). Accordingly, the impact of SPHK inhibitors on translation is alleviated by the concomitant inhibition of ISR kinases. On the other hand, and despite the large number of molecules tested, our study failed to identify chemicals capable of substantially increasing mRNA translation, raising doubts on to what extent translation can be supra-physiologically stimulated in mammalian cells. In summary, our study provides the first comprehensive characterization of the effect of known drugs on protein translation and has helped to unravel a new link between lipid metabolism and mRNA translation in human cells.

## INTRODUCTION

mRNA translation is a fundamental step in gene expression and constitutes the most energy-demanding process in living cells (1, 2). Accordingly, protein synthesis rates are very tightly regulated and rapidly adapt in response to stimuli through the activation of various signaling pathways. For instance, activation of the mammalian target of rapamycin complex 1 (mTORC1) pathway in response to nutrients or growth factors leads to stimulation of protein synthesis (3). In contrast, different sources of stress such as amino acid or heme deprivation, viral infection and endoplasmic reticulum (ER) stress trigger the so-called integrated stress response (ISR), which, in contrast to mTORC1, lowers global rates of protein synthesis (4). The relevance for proper translational regulation is evidenced by the fact that alterations in translation have been associated to a wide variety of human diseases, the number of which has recently increased due to advances in genome-wide association studies (reviewed in (5, 6)). Cancer, immunodeficiency, metabolic and neurological disorders are some examples of diseases linked to aberrant protein synthesis. In addition, low levels of translation have been linked to several age-related degenerative conditions.

In what regards to cancer, high cell proliferation rates demand a corresponding increase in the biosynthesis of cellular components, and several observations suggest a contribution of increased translation levels to carcinogenesis. For instance, a number of oncogenes drive transcription of components of the translation machinery and the over-expression of translation initiation factors can facilitate oncogenic transformation (reviewed in (7, 8)). In this context, several strategies have focused on lowering translation levels in cancer cells for oncological therapy (9). For example, chemical or genetic inhibition of the eIF4F complex involved in translational initiation has shown promising results in overcoming the resistance to various cancer therapies, and several compounds are in clinical trials as antineoplastic drugs (10). An indirect strategy to lower translation is based on small molecules that inhibit rRNA transcription, and compounds such as Actinomycin D (Act D) are currently approved for their use in cancer chemotherapy. Besides drugs that target translation, most therapeutic efforts rely on targeting the signaling pathways that regulate protein metabolism. The best example is that of compounds targeting mTORC1, which are already approved for medical use in oncology and to reduce host versus graft-rejection in organ transplants. Another alternative to lower translation rates in cancer cells is through the use of chemicals that activate the ISR (4). In contrast, inactivating the ISR with chemical inhibitors of the PERK kinase such as ISRIB has shown particularly promising results in the context of neurodegenerative diseases (6, 11, 12).

Until recently, measuring overall levels of translation relied on technologies such as analysing the incorporation of radioactive amino acids, which had limited throughput. In fact, only two chemical screening campaigns directed towards the identification of modifiers of translation have been ever reported (13, 14). However, both of them were based on measuring the translation of a specific fluorescent reporter, and no screen has been yet conducted focused on the analysis of overall translation levels. The recent development of non-canonical amino acids such as puromycin derivatives that can be modified by fluorophores or tags to allow the detection of newly synthesized proteins has now revolutionized the field (15). These techniques have improved translation analyses by methods such as mass-spectrometry, ribosome-sequencing or immunofluorescence (16). Here, we have capitalized on this technology and conducted a chemical screen to evaluate how 1,200 medically approved compounds, as well as around 3,000 compounds at an advanced level of development, modulate overall levels of translation in mammalian cells.

## RESULTS

### A chemical screen for modulators of protein synthesis

In order to monitor changes in global protein synthesis we used a labelling method based on o-propargyl-puromycin (OPP) (15). OPP is an analog of puromycin that is incorporated into the C-terminus of newly synthesized peptides and which can be conjugated to fluorophores by click chemistry, enabling its visualization by microscopy. First, to determine the suitability of this assay for a chemical screen, we evaluated OPP incorporation in 384-well plates using human osteosarcoma-derived U2OS cells. Image-analysis using High-Throughput Microscopy (HTM) revealed a generalized incorporation of OPP in control wells, which was significantly reduced by the use of the mTOR inhibitor Torin 2 (0.5 μM, 3h), and virtually disappeared after treatment with the translation elongation inhibitor cycloheximide (CHX; 100 μg/ml, 30 min) (**Fig. 1 A**,**B**). Of note, while a large fraction of OPP signal is located in the cytoplasm, we also detected a small fraction in the nucleus, which has been previously reported (17). Nevertheless, as translation takes places in the cytoplasm, we used the quantification of the cytoplasmic OPP signal for subsequent analyses.

**Figure 1.**
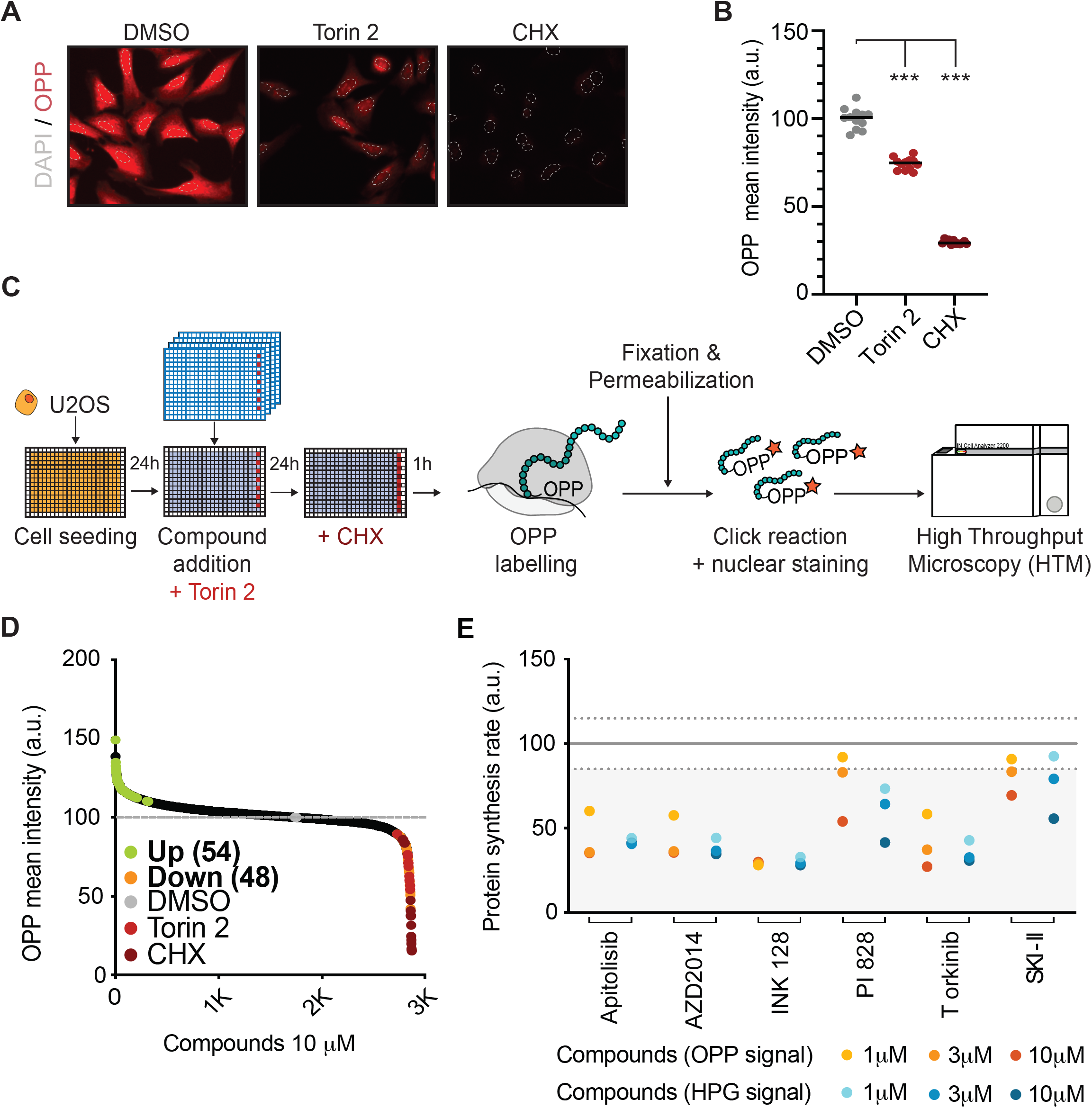
A Chemical Screen for modulators of protein synthesis. (A) OPP incorporation (RED) in U2OS cells exposed to the mTORC1 inhibitor Torin 2 (0.5 μM, 3h) or cycloheximide (CHX, 100 μg/mL, 30 min). Nuclei were stained with Hoechst (dashed GREY line). (B) High-throughput microscopy (HTM)-mediated quantification of OPP-associated cytoplasmic signal in U2OS cells exposed to Torin 2 and CHX, as in (A). ***p < 0.001. (C) Schematic overview of the phenotypic screen workflow. U2OS cells were plated in 384-well plates. On the next day, cells were exposed to compounds at 10μM or 0.5 μM of Torin 2 in specific wells (RED) as a control. After 23h, CHX (100 μg/mL) was added for an hour in specific wells as an additional control (DARK RED). Then, all wells were pulsed with OPP for an hour, after which cells were fixed and processed for HTM-dependent quantification of the OPP signal and nuclei counts. (D) Compound distribution from the screen described above. Compounds increasing (up-regulators, GREEN) or decreasing (down-regulators, ORANGE) OPP incorporation over 3 standard deviations compared to the DMSO control (GREY) are shown. Compounds exceeding 30% toxicity were excluded from this analysis. (E) Dose response (1, 3, 10 μM) of 6 down-regulators after exposing U2OS to the compounds for 24h. Hit validation was conducted by measuring OPP-(ORANGE) or HPG-(BLUE) incorporation assays.

Next, using this setup we performed a chemical screen to evaluate how 4,166 pharmacologically active annotated compounds, including approximately 1,200 that are medically approved, modulate OPP incorporation in U2OS cells. The screen was conducted in triplicate plates, in which each compound was present at 10 μM for 24h (**FIG 1C**). After HTM-mediated evaluation of OPP intensity, we represented the distribution of the effects of all compounds on translation levels (**Fig. 1D; Table S1**). As expected, Torin 2 and CHX were consistently found among the compounds that reduced the OPP signal, with no compound lowering it beyond the effect observed with CHX. For subsequent validation experiments, we selected as hits compounds that met 4 criteria: (a) an increase or decrease in OPP intensity greater than 3 standard deviations over the average signal of DMSO controls; (b) an effect in cellular viability not greater than 30%; (c) that the compound appeared as a hit in all 3 triplicate plates, and (d); that the variation coefficient between the 3 triplicate plates was lower than 20% for both OPP intensity and nuclear counts (viability). With these constraints, 54 compounds that increased OPP incorporation (up-regulators) and 48 compounds that decreased it (down-regulators) were selected for validation experiments. Noteworthy, while most down-regulators were annotated as mTOR/PI3K/MAPK inhibitors, there was no enrichment for a specific chemical group among the up-regulators.

### Validation screens failed to identify chemicals increasing translation

Next, we conducted a secondary validation screen with the selected hits, although we reduced the representation of mTOR/PI3K/MAPK inhibitors as the effect of this class of compounds in translation is well established. For validation, each compound was tested in two independent plates at 3 concentrations (1, 3, 10 μM) for 24h, and the effects in translation were evaluated in two orthogonal assays: (1) quantifying OPP incorporation and (2); measuring the incorporation of homopropargylglycine (HPG), an analogue of methionine that, in contrast to OPP, does not arrest translation after its incorporation into elongating polypeptides (18). While these analyses showed mild effects for seven of the up-regulators (**FIG S1A; Table S2**), subsequent experiments presented significant variability and inconsistent effects of these compounds in stimulating OPP incorporation. Moreover, none of the compounds affected OPP incorporation rates in a shorter treatment (3h), suggesting that, if any, the effects observed in translation after a 24h treatment would be indirect (**FIG S1B**). In this context, and as any increase in translation that we ever detected while conducting these experiments was small, we decided to evaluate the feasibility of our image-based approach to detect a substantial effect in stimulating translation. To do so, we exposed U2OS cells to L-Leucine and insulin, which have been broadly used as stimulators of mTORC1 signaling, particularly in cells that have been previously starved from nutrients (19). Activation of mTORC1 in these conditions was confirmed by an increase in ribosomal protein S6 kinase β-1 (P70S6K) phosphorylation, which was prominent in starved cells exposed to insulin, although subtle in response to L-Leucine (**FIG S1C**). However, while, as expected, starvation reduced overall translation rates in U2OS cells, neither addition of L-Leucine or insulin had a significant effect in increasing OPP signal (**FIG S1D**). These results raised doubts on to what extent the image-based OPP-incorporation assay had a sufficient window to detect compounds capable of increasing overall translation levels and drove us to focus on the down-regulators.

### The sphingosine kinase inhibitor SKI-II lowers overall translation levels

In contrast to the discrepant data among translation up-regulators, the dose-response validation experiments mentioned above identified six compounds that consistently decreased translation rates in U2OS cells in a dose-dependent manner as measured by either OPP or HPG incorporation (**FIG 1E**). With the exception of SKI-II, which is an inhibitor of sphingosine kinases (SPHKs) (20), all validated compounds were annotated as mTOR/PI3K/MAPK inhibitors. We thus focused on further characterizing the effects of SKI-II in translation. First, and in order to test whether the effects of SKI-II were direct, we exposed cells to the compound for 3h or 24h. SKI-II significantly decreased OPP incorporation at both time points, although to a lesser extent than Torin 2 (**FIG 2 A, B**). Next, we evaluated the impact of SKI-II treatment in the activity of the two main signaling cascades controlling translation, mTORC1 and the ISR. Unlike Torin 2, treatment with SKI-II did not affect the phosphorylation of mTORC1 targets such as P70S6K or the eukaryotic translation initiation factor 4E-binding protein 1 (4E-BP1). In contrast, and similar to what is observed with Tunicamycin, a well-established activator of the ISR, SKI-II triggered phosphorylation of the eukaryotic translation initiation factor 2 subunit 1 (eIF2α) and increased the levels of activating transcription factor 4 (ATF4) (**FIG 2C**). Moreover, SKI-II treatment led to the nuclear accumulation of ATF4, which occurs subsequent to eIF2α phosphorylation during activation of the ISR (**FIG 2D**).

**Figure 2.**
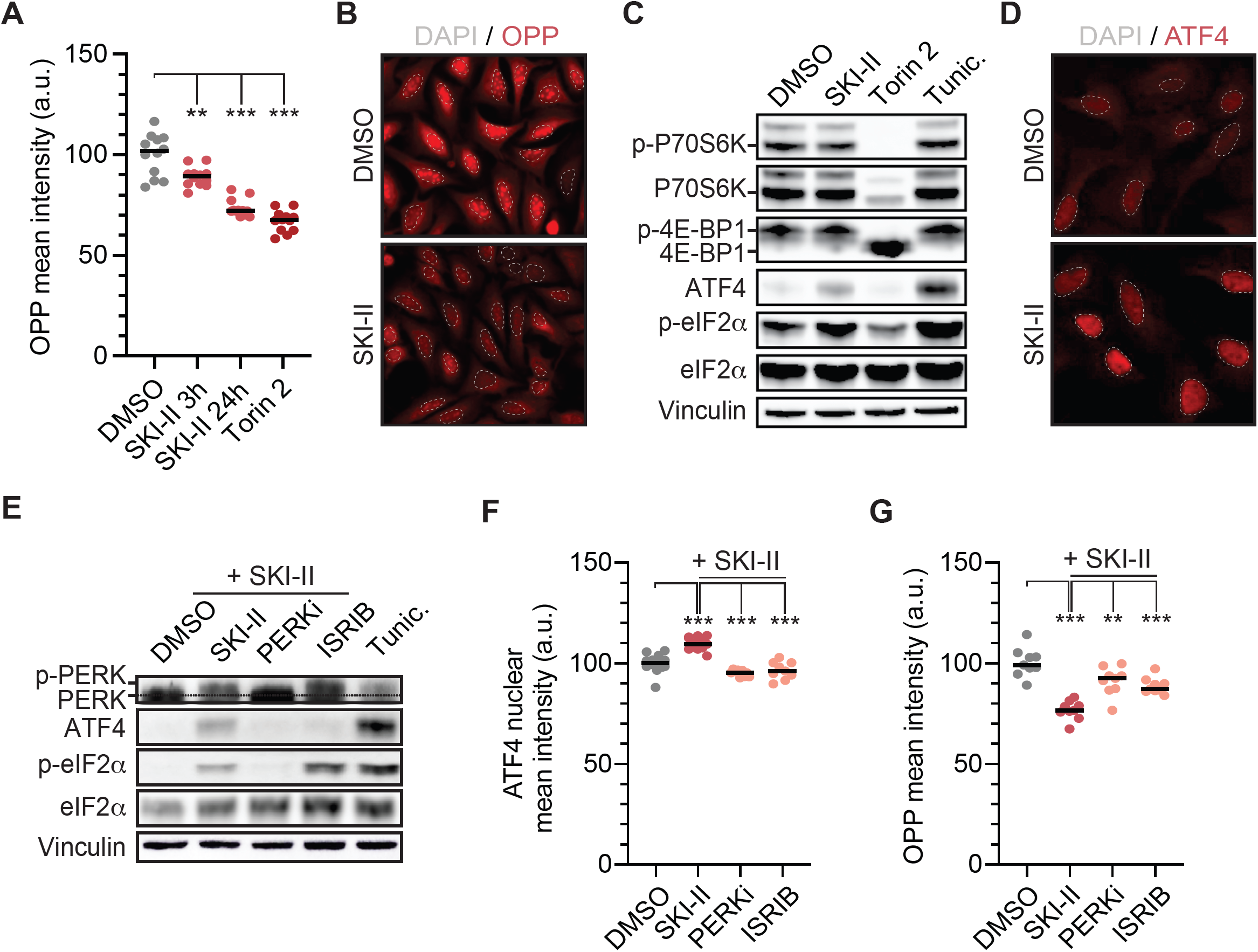
SKI-II reduces protein synthesis by activating the ISR. (A) HTM-mediated quantification of the mean intensity of OPP signal in U2OS cells exposed to SKI-II (10 μM) at different time points (3 and 24h). Torin 2 (0.5 μM, 3h) was included as a control. (B) Reduction in OPP signal (RED) in U2OS cells exposed to SKI-II (10 μM) for 3h compared to the DMSO control. Nuclei were stained with Hoechst (dashed GREY line). (C) Western blot (WB) for markers of the mTORC1 (p-P70S6K, 4E-BP1) or ISR (p-eIF2α, ATF4) pathways in U2OS cells exposed during 3h to SKI-II (10 μM), Torin 2 (0.5 μM). The ISR activating compound Tunicamycin (Tunic., 10 μg/mL) or the mTORC1 inhibitor Torin 2 were used as controls (0.5 μM, 3h). Total levels of P70S6K and eIF2α are shown for normalization. (D) Nuclear accumulation of ATF4 in U2OS cells exposed to SKI-II (10 μM) for 3h. (E) WB of markers of activation of the ISR (p-PERK, ATF4 and p-ei2Fα) in U2OS cells pre-exposed for 1h to PERK inhibitor (PERKi, 0.5 μM) and ISRIB (50 nM) or DMSO, and then exposed to SKI-II (10 μM) for 3h. The phosphorylated form of PERK (p-PERK) is recognized as a shift in the migration pattern of the protein. (F) HTM-mediated quantification of ATF4 mean nuclear intensity in U2OS cells treated as in (E). (G) HTM-mediated quantification of OPP signal in U2OS cells treated as in (E). For the HTM-mediated quantifications, every dot represents the mean value of the measurements taken per well from three independent experiments. Statistical analyses were done using One-way ANOVA tests. ***p < 0.001.

Next, we explored whether ISR inhibitors could revert the effects of SKI-II in translation. In addition to increasing eIF2α phosphorylation and ATF4 levels, exposing cels to SKI-II or Tunicamycin induced phosphorylation of PERK kinase, a critical mediator of the ISR, as evidenced by its reduced mobility in western blotting (WB) (**Fig. 2E**). Moreover, exposure to the PERK inhibitor GSK2606414 (PERKi) (21) blocked PERK activation, phosphorylation of eIF2α and the increase in the levels of ATF4 induced by SKI-II. In contrast, ISRIB, which inhibits the ISR downstream of PERK and eIF2α (22), solely prevented the accumulation of ATF4 but not PERK or eIF2α phosphorylation. Consistent with WB data, HTM-mediated analysis showed that PERKi and ISRIB prevented the nuclear translocation of ATF4 induced by SKI-II (**FIG 2F**). Moreover, PERKi and ISRIB reverted the downregulation of protein synthesis induced by SKI-II, as measured by OPP incorporation (**FIG 2G**). To evaluate the effects of SKI-II on translation in an independent assay, we performed polysome analyses in the breast cancer cell line MCF7. Of note, we chose MCF7 as these cells were more suited for polysome profiling than U2OS, and OPP incorporation experiments revealed a similar effect of SKI-II in reducing translation in both cell types (**Fig. S2A**). Consistent with lower levels of translation, exposure to SKI-II resulted in an accumulation of 80S monosomes in MCF7 cells that was associated to a loss of polysomes, which was reverted by a concomitant treatment with PERKi (**FIG S2B**). Collectively, these data identify SKI-II as a chemical that lowers overall translation levels in human cells, which is mediated by the activation of the ISR.

### SKI-II damages the membranes of the endoplasmic reticulum (ER)

To better understand the mechanism by which SKI-II impacts on translation, we conducted proteomic analysis in U2OS cells exposed to this compound (6h; 10 μM) (**FIG 3A**). SKI-II significantly modified the levels of 43 proteins. Interestingly, some of the most highly up-regulated proteins such as HERPUD, CRE3BL2, XBP1 and DNAJB9, are associated to pathways related to the ISR-activating ER-stress response, including the Unfolded Protein Response (UPR) and ER-associated protein degradation (ERAD) system (**FIG 3A**). Accordingly, Gene Set Enrichment Analysis (GSEA) revealed a significant enrichment of “ER-unfolded protein response” and “negative regulation of response to ER stress” pathways in response to SKI-II (**FIG 3B)**. Thus, the main cellular effects of SKI-II are related to a specific perturbation of the ER, which leads to the activation of the ER-stress pathways.

**Figure 3.**
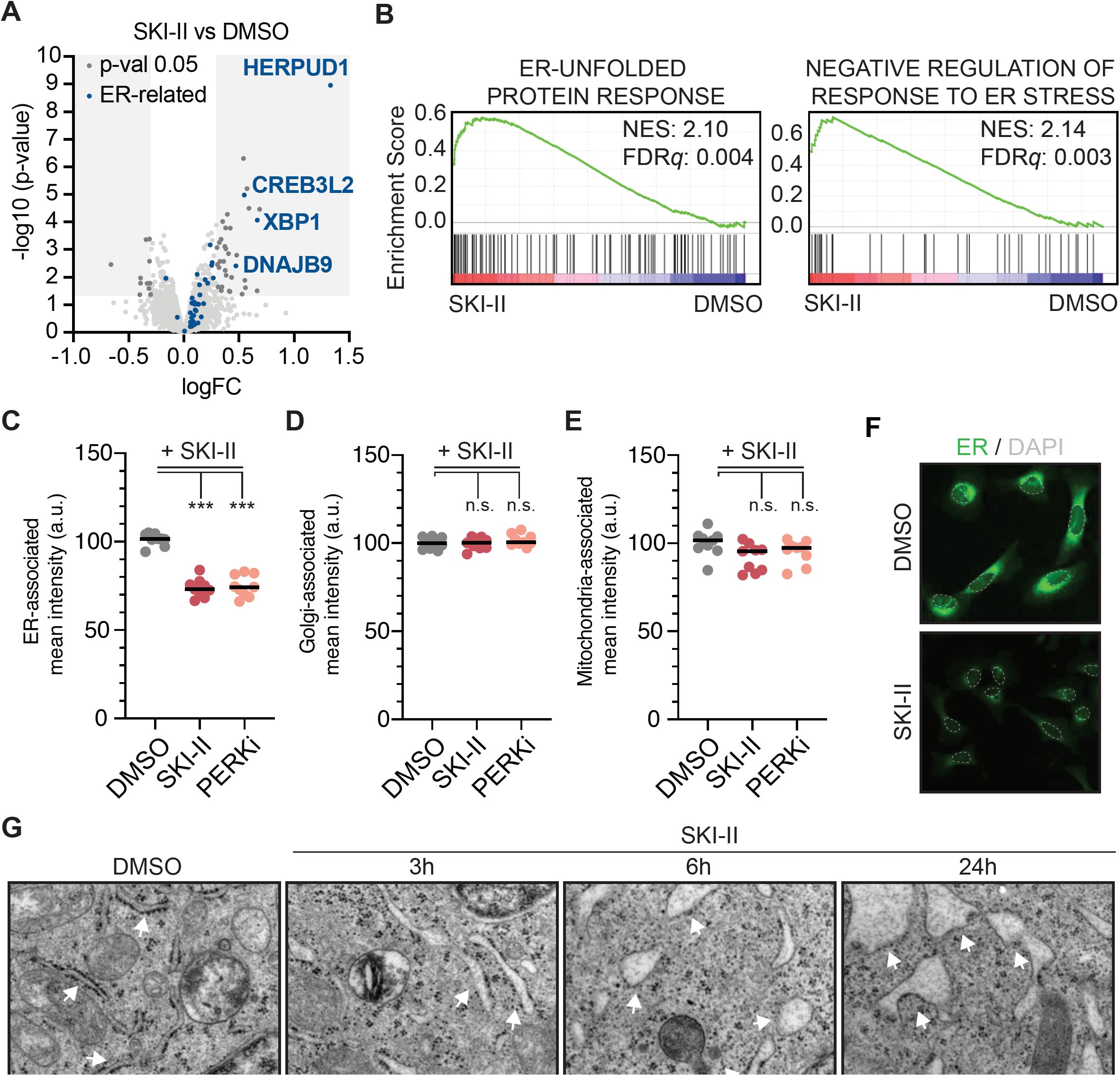
SKI-II specifically affects ER membranes. (A) Volcano plot representing the data from a proteomic experiment in U2OS cells treated with SKI-II (10 μM, 6h) compared to a DMSO-treated control. Proteins within the GREY boxes are differentially regulated (p< 0.05). Factors in BLUE are related to the ER stress response. (B) Gene Set Enrichment Analysis (GSEA) from the experiment defined in (A) showing a significant enrichment in Gene Ontology (GO) terms “ER-unfolded protein response” (NES: 2.10, FDRq: 0.004) and “Negative regulation of response to ER stress” (NES: 2.14, FDRq: 0.003) for cells exposed to SKI-II. (C-E) Quantification from a Cell Painting experiment in U2OS cells treated with SKI-III (10μM, 3 h). ER (C), Golgi (D) and mitochondria (E) were labelled with Concanavalin A, MitoTracker and wheat germ agglutinin, respectively, and signals quantified by HTM. (F) Representative image illustrating the reduction in ER-associated signal (GREEN) in U2OS cells exposed to SKI-II (10 μM) for 3h compared to the DMSO control. Nuclei were stained with Hoechst (dashed GREY line). (G) Transmission Electron Microscopy images of U2OS cells treated with SKI-II (10 μM) for 3, 6 and 24h. Cells were exposed to DMSO as a control. White arrows point at the ER. Ribosomes are black electrodense particles situated around the cisterns of the ER. Scale bars, 0.5 μm. In HTM-quantifications, every dot represents the mean value of the measurements taken by well from three independent experiments. Statistical analyses were done using One-way ANOVA tests. ***p < 0.001.

Given that SKI-II is a sphingosine kinase inhibitor and that sphingolipids and their associated metabolites form part of cellular membranes (23), we evaluated the effect of SKI-II in several membrane-associated organelles. To do so, we used Cell Painting, a technique developed to label cellular organelles for their analysis by fluorescence microscopy (24). HTM-mediated quantification of Cell Painting in U2OS cells treated with SKI-II revealed a significant loss of the ER signal, with no apparent changes in the mean intensity of the signal associated to the Golgi apparatus or mitochondria (**FIG 3C-F**). Noteworthy, the reduction in ER-associated signal in cells treated with SKI-II was not rescued by the inhibition of PERK, arguing that this is a direct effect of the compound on the ER-membrane that would then trigger the activation of the ISR (**FIG 3C**). Supporting this view, Transition Electron Microscopy (TEM) of U2OS cells treated with SKI-II revealed an enlargement of the ER cisterns already evident after 3h, which was exacerbated at times 6h and 24h. This phenomenon was accompanied by a de-attachment of ribosomes from ER membranes, potentially contributing to the overall reduction in protein synthesis triggered by SKI-II. Nevertheless, changes in ER morphology can be produced directly by changes in the cellular lipid composition or as a consequence of an activation of PERK-dependent signaling of the Unfolded Protein Response (UPR) (25, 26). To distinguish between these two possibilities, we compared the effects of SKI-II to those of Thapsigargin, a compound that inactivates Ca^2+^-dependent chaperones necessary for protein folding in the ER and thus activates the UPR (27). While both SKI-II and Thapsigargin reduced the ER signal as measured by Cell Painting to a similar extent, PERK inhibition only rescued the ER signal after treatment with Thapsigargin and had no effect in the context of SKI-II (**FIG S3A**). Together, these results support that the primary effect of SKI-II is a specific damage to ER membranes, which triggers the activation of the ISR thereby reducing translation rates in mammalian cells.

### SPHK2 mediates the toxicity, and ISR activation, triggered by SPHK inhibitors

Besides its effects in translation, whether an activated ISR contributes to cellular toxicity is less explored. In this regard, we observed that inhibition of the ISR with either PERKi or ISRIB rescued the toxicity of Tunicamycin, along with its effects on translation or ATF4 nuclear accumulation (**Fig. S3B**-**E**). Moreover, and similar to SKI-II, Tunicamycin perturbed ER membranes as measured by Cell Painting or TEM (**Fig. S3F-G**). The similarities between these two compounds made us revisit the mechanism of toxicity that has been classically proposed for SPHK inhibitors, which was focused on the accumulation of ceramides as inductors of apoptosis (28). Supporting that the activation of the ISR contributes to the toxicity of SKI-II, nuclear counting experiments or clonogenic survival assays revealed that PERKi or ISRIB reduced the toxicity of SKI-II in U2OS cells (**FIG 4A**,**B**). Similarly, PERKi or ISRIB reversed the effects of SKI-II in cell survival in a mouse cell line of acute myeloid leukaemia (AML), which is relevant given that SKI-II was shown to be particularly efficacious for the killing of AML cells *in vitro* and *in vivo* (29) (**Fig. 4C**).

**Figure 4.**
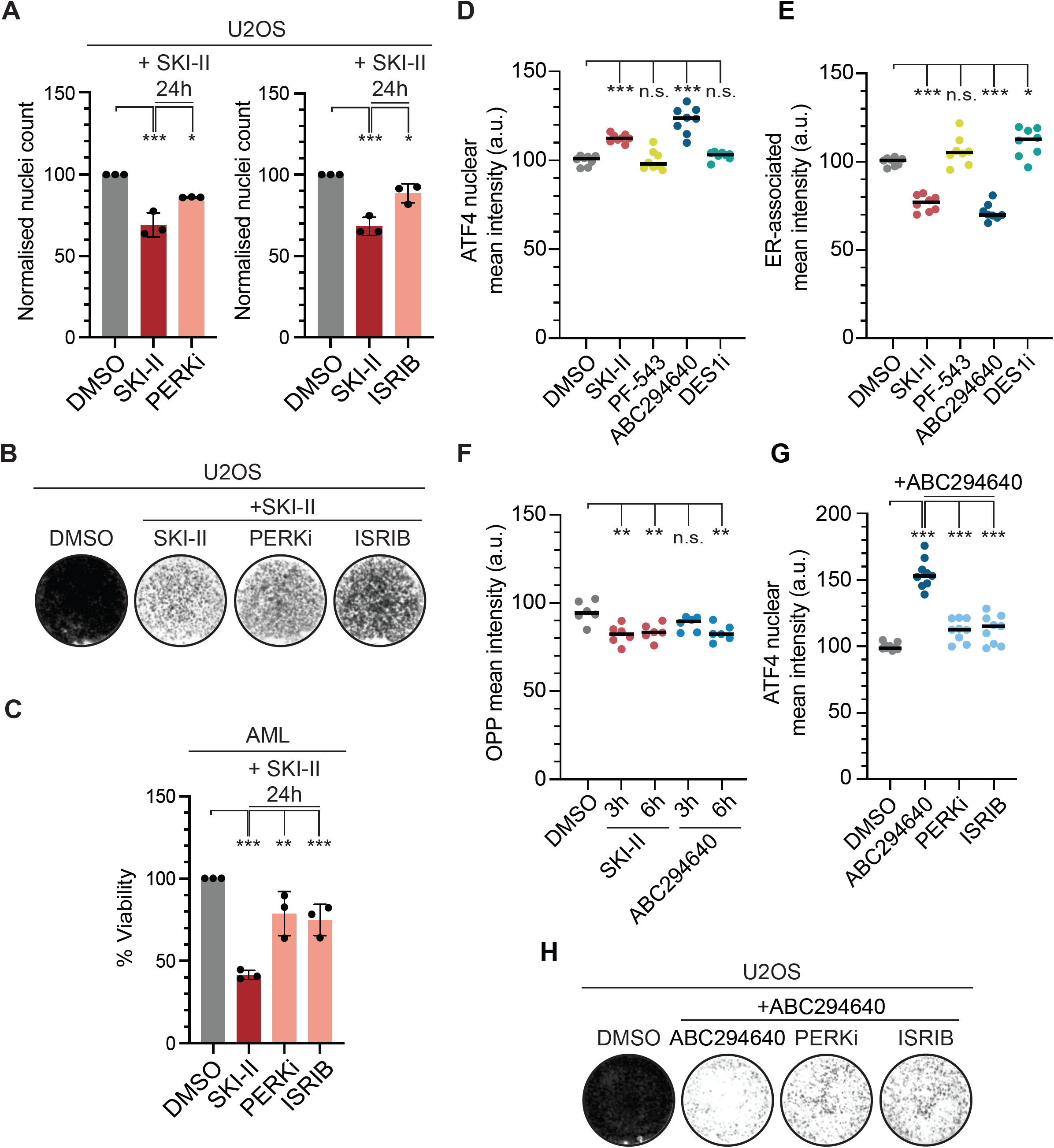
ISR inhibition reduces the toxicity of SKI-II and its clinical derivative ABC294640. (A) HTM-dependent quantification of nuclei numbers (normalized to DMSO) in U2OS cells exposed to SKI-II (10 μM), alone or in combination with PERKi (0.5 μM) or ISRIB (10 nM) for 24h. Statistical analyses were done using One-way ANOVA tests. ***p < 0.001. (B) Clonogenic survival assay of U2OS cells exposed to SKI-II (10 μM), alone or in combination with PERKi (0.5 μM) or with ISRIB (10 nM) for 10 days. (C) Percentage of cellular viability (normalized to DMSO) measured with Cell Titer Glo (CTG) in mouse AML cells exposed to SKI-II (1 μM), alone or in combination with PERKi (3 μM) or ISRIB (10 nM) for 24h. Statistical analyses were done using One-way ANOVA tests. ***p < 0.001. (D) HTM-mediated quantification of ATF4 nuclear signal in U2OS cells treated with SKI-II (10 μM), the SPHK1 inhibitor PF-543 (10 μM), the SPHK2 inhibitor ABC294640 (50 μM) or the DES1 inhibitor DESi (10 μM) for 3h. (E) HTM-mediated quantification of the mean intensity of the ER signal (measured by Cell Painting) in U2OS cells treated as in (D). (F) HTM-quantification of OPP signal in U2OS exposed to SKI-II (10 μM) or ABC294640 (50 μM) for 3 and 6h. (G) HTM-dependent quantification of ATF4 mean nuclear intensity in U2OS pre-exposed for 1h to PERKi (0.5 μM), ISRIB (50 nM) or DMSO, and then exposed to ABC294640 (50 μM) for 3h hours. (H) Clonogenic survival assay of U2OS cells exposed to ABC294640 (50 μM), alone or in combination with PERKi (0.1 μM) or with ISRIB (10 nM) for 10 days. For HTM-mediated quantifications, every dot represents the mean value of the measurements taken by well from three independent experiments. For all cases, statistical analyses were done using One-way ANOVA tests. ***p < 0.001.

SKI-II is a first-generation sphingosine kinase inhibitor, an analogue of sphingosine that inhibits both SPHK1 and SPHK2, with more preference for the latter in biochemical assays (20, 30). Furthermore, SKI-II has been shown to inhibit desaturase 1 (DEGS1), which is also involved in the synthesis of sphingolipids (31). We thus wanted to clarify which of these targets was responsible for the effects of SKI-II in toxicity and translation. To do so we exposed U2OS cells to SKI-II or to selective inhibitors of SPHK1 (PF-543) (32), SPHK2 (ABC294640) (33) and DEGS1 (DESi) (31) for 3h. Interestingly, only the treatment with ABC294640 led to a significant increase in nuclear ATF4 **(FIG 4D)**, a reduction in ER signal measured with Cell Painting **(FIG 4E)** and to lower OPP-incorporation rates (**Fig. 4F**). ABC294640 is a structural analog of SKI-II which was developed to be more selective for SPHK2 than for SPHK1, and is currently in clinical trials for cancer chemotherapy (31). Based on our previous results with SKI-II, and given the clinical relevance of ABC294640, we set to explore whether its mechanism of toxicity was also related to an activation of the ISR. In support of this, PERKi or ISRIB rescued the nuclear accumulation of ATF4 triggered by ABC294640 and reduced its toxicity in clonogenic survival assays (**Fig. 4G-H**). Collectively, these experiments revealed that the toxicity of SPHK inhibitors, including the clinical compound ABC294640, is in part mediated by a role of SPHK2 in safeguarding the integrity of ER membranes and thus preventing the activation of the ISR.

## DISCUSSION

We here present our results on a screening oriented to evaluate the effects of over 4,100 compounds, including approximately 1,200 medically approved drugs, on overall levels of translation. One general conclusion that emerges from our study is that increasing translation beyond supraphysiological levels seems not trivial, which might be related to the fact that translation is the biological process that utilizes the largest fraction of the energy in living cells (1). In fact, while translation inhibitors have been broadly developed (9), there are very few examples described to enhance protein production, always at modest levels, and mostly related to cells where translation levels were already reduced. For instance, while over-expression of the translation factor eIF1A rescues translation in a cellular model of amyotrophic lateral sclerosis where protein synthesis was compromised, it fails to stimulate translation in control conditions (34). Along these lines, expression of an active form of P70S6K or deletion of 4E-BP1, both of which activate mTORC1 signaling, did not result in an accumulation of polysomes in control conditions in *Drosophila* (35). As for the ISR, whereas inhibition of the ISR kinase PKR by two small molecules was found to stimulate translation in rabbit reticulocyte extracts, this was due to a basal activation of the ISR in the *in vitro* system and was not reproduced in unchallenged cells (14). Similarly, our work and that of others show that inhibition of the ISR, using PERKi or ISRIB, does not increase protein synthesis in the absence of stress (22). Arguably, the best-known example of a molecule capable of stimulating translation is insulin. However, insulin treatments are performed in cells undergoing starvation where translation rates are reduced (36). Of note, and while we were able to detect an activation of mTORC1 signaling in starved cells exposed to L-Leucine or insulin, none of these treatments led to a detectable increase in OPP incorporation. While it is possible that the image-based approach that we employed fails to detect subtle increases in translation, an alternative interpretation is that translation cannot be substantially stimulated over physiological levels in mammalian cells.

As for molecules reducing translation rates, the fact that most compounds that significantly lowered translation were annotated as mTOR/PI3K/MAPK inhibitors reinforces the central role of this pathway in protein synthesis. In addition, our screen identified the SPHK inhibitor SKI-II as a novel inhibitor of protein synthesis. Our data are consistent with previous observations of ER-stress in response to SKI-II, particularly in combination with temozolomide or bortezomib (37, 38). Our study is nevertheless first in reporting the effects of SK-II in translation and has also helped to clarify the relative role of the different SPHKs. The interest in SPHKs as anticancer targets was initiated from the fact that these enzymes are frequently overexpressed in tumors (28). The original and still widely accepted hypothesis was that inhibition of SPHKs would result in lower levels of the pro-survival factor sphingosine-1-phospate (S1P) while increasing the levels of the apoptotic factor ceramide. Based on this line of thought, several inhibitors of SPHK1, SPHK2 and DEGS1 have been discovered and are at various stages of development as clinical drugs. However, the role of the S1P/ceramide balance and the contribution of each SPHK to the toxicity of these compounds remains to be fully established.

The initial compounds targeting SPHKs, such as SKI-II, were inactive sphingosine analogues that targeted both SPHK1 and SPHK2. However, while early studies pointed to SPHK1 as the relevant anticancer target, later works casted doubts on such a view (39, 40). For instance, even if SPHK1 or SPHK2 depletion both cause defects in cell growth, these are more prominent when targeting SPHK2, yet only SPHK1 depletion leads to a significant increase in ceramides (41-44). Moreover, and contrary to depletion experiments, use of the SPHK1 selective inhibitor PF-543 fails to induce cell death in different cancer cell lines (45). These results indicate that SPHK2 is the main target responsible for the toxicity of SPHK inhibitors and further raise doubts on whether the S1P/ceramide balance is the main determinant of their toxicity. An alternative model is that this toxicity derives from perturbations in membrane fluidity generated after SPHK2 inhibition through the accumulation of ceramide, which would then activate the UPR and ISR (25, 38). Moreover, while SPHK1 is predominantly cytoplasmic, SPHK2 is mainly associated to the ER and mitochondrial membranes (39), supporting a direct effect of SPHK2 inhibitors in the ER composition. In concordance this view, we here report that SKI-II and the SPHK2 selective inhibitor ABC294640 cause direct alterations to the ER and lower translation through activation of the ISR.

While our work here focused on translation, a better understanding of the mechanisms of toxicity of SPHK inhibitors should help to optimize their use as anticancer agents by designing better drug combinations. For instance, induction of ER stress by lipotoxicity potentiates the anticancer effects of agents that activate the UPR such as proteasomal inhibitors (38, 46). On the other hand, we here show, for the first time, that the toxicity of ISR activators such as Tunicamycin or SKI-II is alleviated by a concomitant treatment with PERKi or ISRIB, although whether this is related to translation remains unknown. To end, and despite ISR activators might be toxic for cancer cells, we should note that there are also disorders where an induction of the ISR has been proposed to be beneficial, such as Charcot-Marie-Tooth disease, opening new areas where SPHK inhibitors might be of help (11, 47). In addition, mTORC1 inhibitors that lower translation are widely used in biomedical research as tools to revert age-related pathologies and some have even obtained clinical approval in the context of oncology and organ transplant. To what extent other drugs that lower overall translation levels independently of mTORC1 might be of use in these or other pathologies emerges as an interesting possibility.

## MATERIALS AND METHODS

### Cell lines

Human female osteosarcoma U2OS and breast cancer MCF7 cells were cultured in DMEM + Glutamax (Thermo Fisher Scientific, 31966-047) supplemented with 10% FCS and 1% Penicillin/Streptomycin. Mouse AML cells (48), were cultured in RPMI1640 (Thermo Fisher Scientific, 3196621875034), also supplemented with serum and antibiotics. For the validation screen, DMEM lacking methionine (Thermo Fisher Scientific, 21013024) and supplemented with 2mM L-glutamine (Sigma-Aldrich, G7513) and 5mM L-Cysteine (Sigma-Aldrich, C6727) was used. The identity of the cell lines with human origin, U2OS and MCF7, was confirmed by short tandem repeat (STR) profiling analysis (STRA11385, STRA11386).

### Compounds

Torin 2 (SML1224), cycloheximide (CHX, C7698), insulin solution (I0516), L-Leucine (L8000), Clomiphene citrate (C6272), ARC239 dihydrochloride (A5736), Imatinib (SML1027), UNC1999 (SML0778), SKI-II (S5696), and ISRIB (SML0843) were purchased from Sigma-Aldrich; NVP-BVU972 (S2761), FK-506 (Tacrolimus, S5003) were obtained from Selleck Chemicals, JNK Inhibitor V (420129) from Calbiochem; PERK inhibitor GSK2606414 (PERKi, 516535) from Merck Millipore; Tunicamycin (11445) from Bionordika; Thapsigargin (ab120286) from Abcam; PF-543 hydrochloride (5754) from R&D Systems; ABC294640 (10587) from Cayman chemical and the DES1 inhibitor (B-0027) from Echelon biosciences.

### Chemical Screens

Plate and liquid handling were performed using Echo®550, (Labcyte), Janus® Automated Workstation, (PerkinElmer), Multidrop 384 (Thermo Scientific), Viaflo 384 (Integra Bioscience), Multiflo FX Multi-Mode Dispenser (BioTek) and Hydrospeed™ washer (Tecan). Cells were seeded in black with clear bottom 384-well plates (BD Falcon, 353962). Compound libraries were provided by the Chemical Biology Consortium Sweden (CBCS). The chemical collection used in the primary screening contained 4,166 pharmacologically active compounds from the following libraries: Prestwick Chemical Library of FDA approved compounds, Tocris mini, Selleck tool compounds, Selleck known kinase inhibitors, ENZO tool compounds, SGC bromodomain probes and 115 covalent drugs synthesized by Henriksson M. (Karolinska Institutet, Sweden).

For the primary screen, U2OS cells were trypsinized and resuspended in culture medium. The cell suspension (750 cells in 30 μl/well) was dispensed into 384-well plates and incubated overnight at 37oC in a 5% CO_2_ atmosphere. The following day, cells were exposed to a final concentration of 10 μM of compounds diluted in dimethyl sulfoxide (DMSO). Compound addition was done in triplicate sets of plates. At the same time, the mTORC1 inhibitor Torin 2 (SML1224, Sigma-Aldrich) was added as a positive control in specific wells at a final concentration of 0.5 μM. After 23 hours, cycloheximide (CHX, C7698, Sigma-Aldrich) was added as an additional control for 1h to a final concentration of 100 μg/mL in specific wells. Cells were then pulsed with OPP (Jena Bioscience) dissolved in media at a final concentration of 20μM for 1h at 37°C. After that, plates were washed once with PBS and cells fixed with 100% ice-cold methanol for 20 min at room temperature (RT). After fixation and washing, cells were permeabilized with Triton X-100 (0.1%) for 20 min, RT. Next, cells were incubated with Click reaction cocktail (88 mM Na-phosphate, pH 7, 20 mM CuSO_4_, 10 mM Na-Ascorbate, 2 μM Alexa Azide 647) for 30 min in the dark. Nuclei were stained with 2 μM Hoechst 33342 for 15 min in the dark. Plates were imaged using an IN Cell Analyzer 2200 system with a 10x objective, four images per well were acquired, covering the whole well. Images were analyzed with CellProfiler using a custom-made pipeline for the detection of cytoplasmic OPP signal. All values were normalized to DMSO conditions within each plate. Then, the mean value for each compound in triplicates was calculated, representing a single measurement per compound per set of triplicates. Images were acquired using an IN Cell Analyzer 2200 (GE Healthcare), and quantitative image analyses were run in the open-source software CellProfiler (www.cellprofiler.org) (49). Statistical analysis of high content imagining data was conducted using TIBCO Spotfire and additional statistical analyses were done using Microsoft Excel or Graphpad Prism software.

For the validation screen, U2OS cells were exposed to three concentrations of the selected hits (final concentration 1, 3 and 10 μM) for 24h. Translation rates were evaluated quantifying both OPP and L-Homopropargylglycine (HPG) incorporation. The validation with OPP and HPG was conducted in duplicates. Prior to the HPG pulse, cells were washed with PBS and media was exchanged to DMEM lacking methionine for 30 min. HPG diluted in DMEM lacking Met was added to cells to a final concentration of 10 μM. Cells were kept at 37°C for 30 min. For HPG incorporation measurements, the Click-iT(tm) HPG Alexa Fluor(tm) 488 Protein Synthesis Assay Kit was used (Thermo Fisher Scientific, C10186).

### Immunofluorescence

U2OS cells were seeded and treated as above onto 96-well imaging plates. Then, cells were fixed with or 4% PFA for 15 min and permeabilized with 0.1% Triton X-100 for 10 min at room temperature. After blocking (3% BSA and 0.1% Tween-20 in PBS) for 30 min, the indicated concentration of primary antibodies was applied: ATF4 (D4B8) (1:200, Cell Signaling Technology, 11815S). The following secondary antibodies were added for 1h at room temperature: anti-rabbit IgG-647 (1:500, Life technologies, A-21244), anti-rabbit IgG-488 (1:500, Life technologies, A-11008). Images were acquired with IN Cell. Nuclear translocation of ATF4 was measured by increase of ATF4 mean intensity into the nucleus of cells using Hoechst as a counterstaining. Cell painting (24) was used to differentially stain membrane-bound organelles. Before fixation, MitoTracker Deep Red A-647 (Thermo Fisher Scientific, M22426) was diluted in DMEM (1:1000) and added to live cells for 20 min, at 37°C. Afterwards, cells were washed with PBS, fixed and permeabilized as before, and stained with a cocktail of Hoechst (1:1000), Concanavalin A A-488 (1:200, Thermo Fisher Scientific, C11252) and Wheat Germ Agglutinin A-555 (1:500, Thermo Fisher Scientific, W32464) for 20 min in the dark. Image analysis was performed using a self-made pipeline identifying the different organelles according to intensity of the signal in the cytoplasm. In these experiments, 9 pictures were taken per well using a 20x objective.

### Western Blotting

RIPA buffer with protease inhibitor (Sigma) was used for preparing protein lysates from U2OS cells treated as indicated. Immunoblotting was performed following standard protocols with indicated antibodies: ATF4 (D4B8) (1:500, Cell Signaling Technology, 11815S), p-P70S6K (Thr389) (1:1000, Cell Signaling Technology, 9205), P70S6K (1:1000, Cell Signaling Technology, 9202S), 4E-BP1 (1:1000, CST Cell Signaling Tech, 9452S), p-eIF2a (phospho S51) [E90] (1:1000, Abcam, ab32157), eIF2a (1:500, Cell Signaling Technology, 9722), Vinculin (EPR8185) (1:2000, Abcam, ab129002), PERK (D11A8) (1:500, Cell Signaling Technology, 5683). Protein bands were visualized by chemiluminescence (ECL, Thermo Scientific, 34076) and imaged on an Amersham Imager 600 (GE healthcare).

### Viability experiments

U2OS (2,000 cells/well) and Mouse AML cells (10,000 cells/well) were seeded onto 96-well plates. The next day, cells were exposed to compounds at indicated concentrations and cultured at 37°C for 24h. Next, U2OS cells were fixed in 4% PFA and nuclei were stained using Hoechst. Images were acquired at 4X magnification with ImageXpress Pico System (Molecular Devices). Nuclei count was done using automatic segmentation using CellReporterXpress (Molecular Devices). As AML cells grow in suspension, CellTiter-Glo Luminescent Assay (CTG, Promega, G7571) was used to assess viability. For clonogenic survival assays, U2OS cells were plated onto 12-well plates (7,000 cells/well). The following day, cells were exposed to compounds at the indicated concentrations. Every other day, the medium was replaced with fresh medium containing compounds until day 10. At the end of the experiment, cells were fixed and stained with 0.4% methylene blue in methanol for 30 min and images acquired with an image analyzer (Amersham Imager 600, GE healthcare).

### Polysome profiling

MCF7 cells (7×10^6^ cells/plate) were seeded in 15cm petri dishes. After 24h, cells were treated as indicated for 20h. Cells were washed and harvested in PBS containing CHX (100 μg/ml). After centrifugation, cells were resuspended in lysis buffer ((5 mM Tris-HCl (pH 7.5), 2.5 mM MgCl2, 1.5 mM KCl and 1x protease inhibitor cocktail (EDTA-free), CHX (100 μg/ml), DTT (2 μM), and 100 units of RNAse inhibitor) followed by addition of Triton X-100 (0.5%) and Sodium Deoxycholate (0.5%), as indicated in (36). Nuclei were removed by centrifugation (14,000 x g for 10 min at 4 °C). Then, lysates were loaded onto sucrose density gradients (15-50% in 20 mM HEPES (pH 7.6), 0.1 mM KCl, 5 mM MgCl_2_, CHX (10 μg/ml), 0.1x protease inhibitor cocktail (EDTA-free), 10 units of RNAse inhibitor). After ultracentrifugation (3h 10min, 37,000 rpm at 4°C using a using SW41Ti rotor), gradients were analysed in a piston gradient fractionator (Biocompo). Profiles were acquired with Gradient profiler v.2.0 (Biocomp) and represented using Graphpad Prism.

### Transmission Electron Microscopy

For the TEM analyses, U2OS (1×10^6^ cells) were plated on 15 cm petri dishes. The following day, cells were fixed at RT in 2 % glutaraldehyde in 0.1 M phosphate buffer, pH 7.4. After fixation, the cells were rinsed in 0.1 M phosphate buffer and centrifuged. Cell pellets were post-fixed in 2% osmium tetroxide in 0.1 M phosphate buffer, pH 7.4 at 4 °C for 2 h. Cells were then stepwise dehydrated in ethanol, followed by acetone and finally embedded in LX-112. Ultrathin sections (∼50–60 nm) were prepared using a Leica EM UC7 and contrasted with uranyl acetate followed by lead citrate. TEM imaging was done in a Tecnai 12 Spirit Bio TWIN transmission electron microscope operated at 80 kV and digital images acquired using a Veleta camera (Olympus Soft Imaging Solutions).

### Mass Spectrometry

For the MS experiments, U2OS cells (1×10^6^) were seeded in 10 cm petri dishes. The next day, cells were treated with SKI-II (10 μM) or DMSO for 6h. Each treatment was done in triplicate. Samples were digested with trypsin using S-trap devices. Peptides were subsequently labeled with TMT-11plex reagents and pre-fractionated by means of High pH-reverse phase chromatography. Fractions were finally analyzed by LC-MS/MS analysis using a Q Exactive Plus (Thermo Fisher Scientific) coupled to an Ultimate 3000 nanoHPLC system. Raw files were analyzed with MaxQuant against a human protein database. Differential analysis was done with ProStar (v1.14) (50): proteins with p.val <0.05 and log2FC >0.3 (<-0.3) were defined as regulated (FDR 5-10% was estimated by Benjamini-Hochberg). Gene Set Enrichment Analysis (GSEA) (v 4.0.2) was performed using the preRanked function (enrichment statistic = weighted). Log2 ratios from each of the comparisons were used to rank all the proteins detected. Gene sets were extracted from Gene Ontology using the Molecular Signature Database (MSigDB) and pathways with a FDR q.value < 0,05 were considered as statistically enriched.

### Data availability

Mass Spectrometry data associated to this work are available via ProteomeXchange with identifier PXD017445.

## ACKNOWLEDGEMENTS

The authors want to thank the Laboratories for Chemical Biology at Karolinska Institutet (LCBKI) for their help with the chemical screening, the Electron microscopy (EMil) Unit at Karolinska Institutet for TEM analyses, Ola Larsson for help with polysome profiling, Jamie Espinoza and Andrea Björkman for discussions, and Kirsten Tschapalda for help in the screening analysis. Work related to this work was funded by grants from the Cancerfonden foundation (CAN 2018/381) and the Swedish Research Council (VR) (538-2014-31) to OF.

## AUTHOR CONTRIBUTIONS

A.C. contributed to most experiments, data analyses and writing of the MS. D.K. contributed to polysome analyses. V.L. contributed to proteomic experiments. M.H. contributed to the chemical screen. J.B. contributed to the MS revision and supervision of polysome experiments. O.F-C. and J.C-P. supervised the study and wrote the MS.

## REFERENCES

1. F. Buttgereit, M. D. Brand, A hierarchy of ATP-consuming processes in mammalian cells. Biochem J 312 (Pt 1), 163–167 (1995).

2. N. Sonenberg, A. G. Hinnebusch, Regulation of translation initiation in eukaryotes: mechanisms and biological targets. Cell 136, 731–745 (2009).

3. G. Y. Liu, D. M. Sabatini, mTOR at the nexus of nutrition, growth, ageing and disease. Nat Rev Mol Cell Biol 21, 183–203 (2020).

4. K. Pakos-Zebrucka et al., The integrated stress response. EMBO Rep 17, 1374–1395 (2016).

5. G. C. Scheper, M. S. van der Knaap, C. G. Proud, Translation matters: protein synthesis defects in inherited disease. Nat Rev Genet 8, 711–723 (2007).

6. S. Tahmasebi, A. Khoutorsky, M. B. Mathews, N. Sonenberg, Translation deregulation in human disease. Nat Rev Mol Cell Biol 19, 791–807 (2018).

7. N. Robichaud, N. Sonenberg, D. Ruggero, R. J. Schneider, Translational Control in Cancer. Cold Spring Harb Perspect Biol 11 (2019).

8. D. Silvera, S. C. Formenti, R. J. Schneider, Translational control in cancer. Nat Rev Cancer 10, 254–266 (2010).

9. J. Chu, J. Pelletier, Therapeutic Opportunities in Eukaryotic Translation. Cold Spring Harb Perspect Biol 10 (2018).

10. M. Bhat et al., Targeting the translation machinery in cancer. Nat Rev Drug Discov 14, 261–278 (2015).

11. V. Gonzalez-Teuber et al., Small Molecules to Improve ER Proteostasis in Disease. Trends Pharmacol Sci 40, 684–695 (2019).

12. S. L. Moon, N. Sonenberg, R. Parker, Neuronal Regulation of eIF2alpha Function in Health and Neurological Disorders. Trends Mol Med 24, 575–589 (2018).

13. O. Novac, A. S. Guenier, J. Pelletier, Inhibitors of protein synthesis identified by a high throughput multiplexed translation screen. Nucleic Acids Res 32, 902–915 (2004).

14. U. Shin et al., Stimulators of translation identified during a small molecule screening campaign. Anal Biochem 447, 6–14 (2014).

15. J. Liu, Y. Xu, D. Stoleru, A. Salic, Imaging protein synthesis in cells and tissues with an alkyne analog of puromycin. Proc Natl Acad Sci U S A 109, 413–418 (2012).

16. S. Iwasaki, N. T. Ingolia, The Growing Toolbox for Protein Synthesis Studies. Trends Biochem Sci 42, 612–624 (2017).

17. N. Uozumi, H. Matsumoto, H. Saitoh, Detection of O-propargyl-puromycin with SUMO and ubiquitin by click chemistry at PML-nuclear bodies during abortive proteasome activities. Biochem Biophys Res Commun 474, 247–251 (2016).

18. D. C. Dieterich, A. J. Link, J. Graumann, D. A. Tirrell, E. M. Schuman, Selective identification of newly synthesized proteins in mammalian cells using bioorthogonal noncanonical amino acid tagging (BONCAT). Proc Natl Acad Sci U S A 103, 9482–9487 (2006).

19. P. Gran, D. Cameron-Smith, The actions of exogenous leucine on mTOR signalling and amino acid transporters in human myotubes. BMC Physiol 11, 10 (2011).

20. K. J. French et al., Discovery and evaluation of inhibitors of human sphingosine kinase. Cancer Res 63, 5962–5969 (2003).

21. J. M. Axten et al., Discovery of 7-methyl-5-(1-{[3-(trifluoromethyl)phenyl]acetyl}-2,3-dihydro-1H-indol-5-yl)-7H-p yrrolo[2,3-d]pyrimidin-4-amine (GSK2606414), a potent and selective first-in-class inhibitor of protein kinase R (PKR)-like endoplasmic reticulum kinase (PERK). J Med Chem 55, 7193–7207 (2012).

22. C. Sidrauski et al., Pharmacological brake-release of mRNA translation enhances cognitive memory. Elife 2, e00498 (2013).

23. D. K. Breslow, Sphingolipid homeostasis in the endoplasmic reticulum and beyond. Cold Spring Harb Perspect Biol 5, a013326 (2013).

24. M. A. Bray et al., Cell Painting, a high-content image-based assay for morphological profiling using multiplexed fluorescent dyes. Nat Protoc 11, 1757–1774 (2016).

25. M. K. Bennett, C. T. Wallington-Beddoe, S. M. Pitson, Sphingolipids and the unfolded protein response. Biochim Biophys Acta Mol Cell Biol Lipids 1864, 1483–1494 (2019).

26. R. Volmer, K. van der Ploeg, D. Ron, Membrane lipid saturation activates endoplasmic reticulum unfolded protein response transducers through their transmembrane domains. Proc Natl Acad Sci U S A 110, 4628–4633 (2013).

27. A. Abdullahi, M. Stanojcic, A. Parousis, D. Patsouris, M. G. Jeschke, Modeling Acute ER Stress in Vivo and in Vitro. Shock 47, 506–513 (2017).

28. B. Ogretmen, Sphingolipid metabolism in cancer signalling and therapy. Nat Rev Cancer 18, 33–50 (2018).

29. L. Yang et al., SphK1 inhibitor II (SKI-II) inhibits acute myelogenous leukemia cell growth in vitro and in vivo. Biochem Biophys Res Commun 460, 903–908 (2015).

30. K. J. French et al., Antitumor activity of sphingosine kinase inhibitors. J Pharmacol Exp Ther 318, 596–603 (2006).

31. L. Aurelio et al., From Sphingosine Kinase to Dihydroceramide Desaturase: A Structure-Activity Relationship (SAR) Study of the Enzyme Inhibitory and Anticancer Activity of 4-((4-(4-Chlorophenyl)thiazol-2-yl)amino)phenol (SKI-II). J Med Chem 59, 965–984 (2016).

32. J. Wang, S. Knapp, N. J. Pyne, S. Pyne, J. M. Elkins, Crystal Structure of Sphingosine Kinase 1 with PF-543. ACS medicinal chemistry letters 5, 1329–1333 (2014).

33. K. J. French et al., Pharmacology and antitumor activity of ABC294640, a selective inhibitor of sphingosine kinase-2. J Pharmacol Exp Ther 333, 129–139 (2010).

34. T. G. Moens et al., C9orf72 arginine-rich dipeptide proteins interact with ribosomal proteins in vivo to induce a toxic translational arrest that is rescued by eIF1A. Acta Neuropathol 137, 487–500 (2019).

35. S. Nagarajan, S. S. Grewal, An investigation of nutrient-dependent mRNA translation in Drosophila larvae. Biol Open 3, 1020–1031 (2014).

36. V. Gandin et al., Polysome fractionation and analysis of mammalian translatomes on a genome-wide scale. J Vis Exp 10.3791/51455 (2014).

37. J. Noack, J. Choi, K. Richter, A. Kopp-Schneider, A. Regnier-Vigouroux, A sphingosine kinase inhibitor combined with temozolomide induces glioblastoma cell death through accumulation of dihydrosphingosine and dihydroceramide, endoplasmic reticulum stress and autophagy. Cell Death Dis 5, e1425 (2014).

38. C. T. Wallington-Beddoe et al., Sphingosine kinase 2 inhibition synergises with bortezomib to target myeloma by enhancing endoplasmic reticulum stress. Oncotarget 8, 43602–43616 (2017).

39. M. R. Pitman, M. Costabile, S. M. Pitson, Recent advances in the development of sphingosine kinase inhibitors. Cell Signal 28, 1349–1363 (2016).

40. W. L. Santos, K. R. Lynch, Drugging sphingosine kinases. ACS Chem Biol 10, 225–233 (2015).

41. P. Gao, C. D. Smith, Ablation of sphingosine kinase-2 inhibits tumor cell proliferation and migration. Mol Cancer Res 9, 1509–1519 (2011).

42. J. R. Van Brocklyn et al., Sphingosine kinase-1 expression correlates with poor survival of patients with glioblastoma multiforme: roles of sphingosine kinase isoforms in growth of glioblastoma cell lines. J Neuropathol Exp Neurol 64, 695–705 (2005).

43. J. K. Venkata et al., Inhibition of sphingosine kinase 2 downregulates the expression of c-Myc and Mcl-1 and induces apoptosis in multiple myeloma. Blood 124, 1915–1925 (2014).

44. C. T. Wallington-Beddoe et al., Sphingosine kinase 2 promotes acute lymphoblastic leukemia by enhancing MYC expression. Cancer Res 74, 2803–2815 (2014).

45. M. E. Schnute et al., Modulation of cellular S1P levels with a novel, potent and specific inhibitor of sphingosine kinase-1. Biochem J 444, 79–88 (2012).

46. C. S. Lewis, C. Voelkel-Johnson, C. D. Smith, Targeting Sphingosine Kinases for the Treatment of Cancer. Adv Cancer Res 140, 295–325 (2018).

47. K. M. Green et al., High-throughput screening yields several small-molecule inhibitors of repeat-associated non-AUG translation. J Biol Chem 294, 18624–18638 (2019).

48. I. Morgado-Palacin et al., Targeting the kinase activities of ATR and ATM exhibits antitumoral activity in mouse models of MLL-rearranged AML. Sci Signal 9, ra91 (2016).

49. T. R. Jones et al., CellProfiler Analyst: data exploration and analysis software for complex image-based screens. BMC Bioinformatics 9, 482 (2008).

50. S. Wieczorek et al., DAPAR & ProStaR: software to perform statistical analyses in quantitative discovery proteomics. Bioinformatics 33, 135–136 (2017).

