## Supplementary material for "A chemical screen identifies a link between lipid metabolism and mRNA translation": Figures S1-3

Corman A *et al.* **Fig.S1**

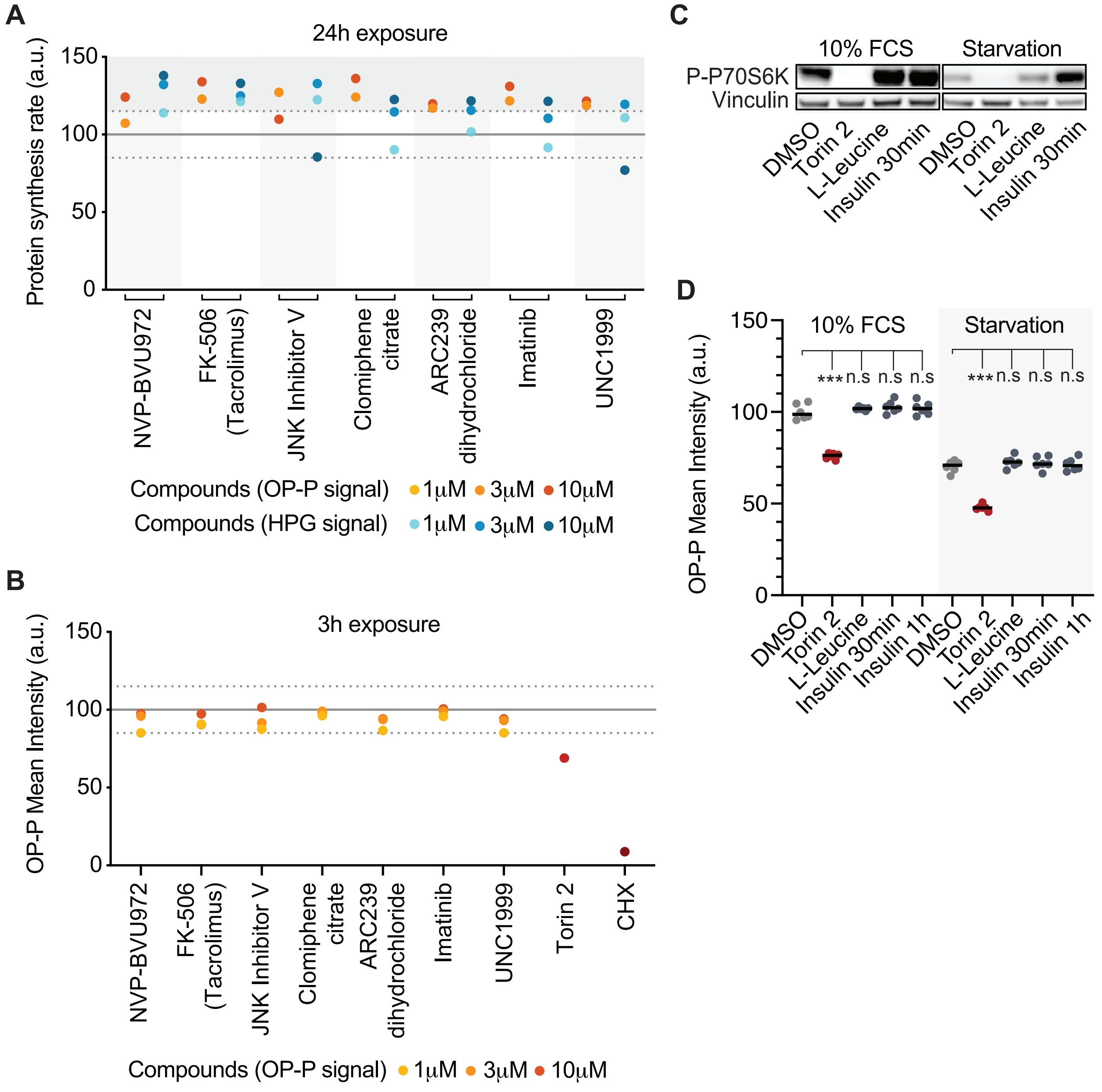

**Figure S1. Characterization of hit compounds stimulating OPP incorporation.** (**A**) Protein synthesis rates (a.u.) of 7 drugs identified as potential up-regulator hits in the primary screen after exposing U2OS to different concentrations (1, 3, 5 μM) of the compounds for 24h. Hit validation was conducted using both OPP (ORANGE) and HPG (BLUE) labelling assays. (**B**) HTM-mediated quantification of OPP signal in U2OS cells exposed for 3h to three concentrations (1, 3, 10 μM) of the compounds annotated as up-regulators of OPP incorporation. Torin 2 (0.5 μM) and CHX (100 μg/mL) were included as controls. DMSO control appears as a GREY line, and a range of 15% appears as a dashed GREY Line. (**C**) HTM-mediated quantification of OPP signal in U2OS cells in the presence of molecules the amino acid L-Leucine (4mM, 4h) and insulin (0.1 μM, 30 min and 1h). Compound treatments were done in cells growing in complete media (10%FCS) and in cells starved over-night (media with no FCS). For the HTM-quantification, every dot represents the mean value of the measurements taken by well from three independent experiments. Statistical analyses were done using One-way ANOVA tests. ***$p < 0.001$. (**D**) WB measuring the phosphorylation of the mTORC1 target P70S6K in cells treated as in (**C**). Vinculin levels are shown for loading control.

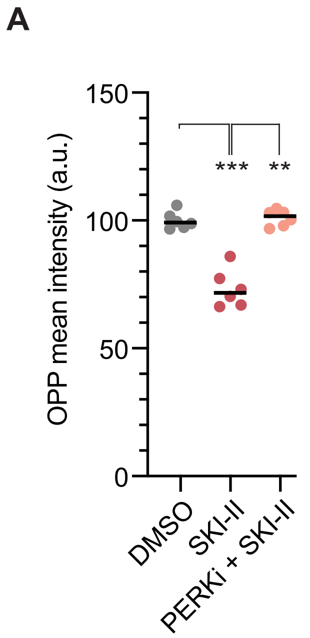
A
150
100
50
0
OPP mean intensity (a.u.)
***
**
DMSO
SKI-II
PERKi + SKI-II

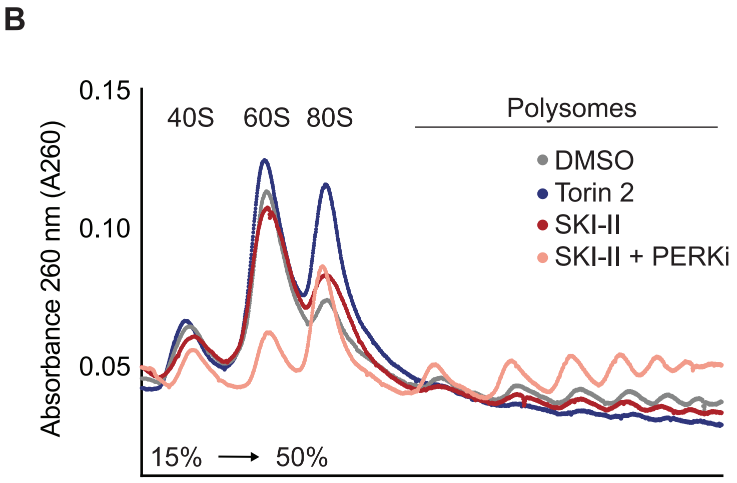
B
0.15
0.10
0.05
Absorbance 260 nm (A260)
40S
60S
80S
Polysomes
DMSO
Torin 2
SKI-II
SKI-II + PERKi
15% → 50%

**Figure S2. SKI-II treatment reduces translation rates in MCF7 cells.** (**A**) HTM-mediated quantification of OPP signal in MCF7 cells exposed to either SKI-II (10 μM), SKI-II with PERKi (0.5 μM), or DMSO for 20h. (**B**) Polysome profiles of MCF7 cells treated as in (A). Torin 2 (0.1 μM) was included as an additional control.

Corman A *et al.* **Fig.S3**

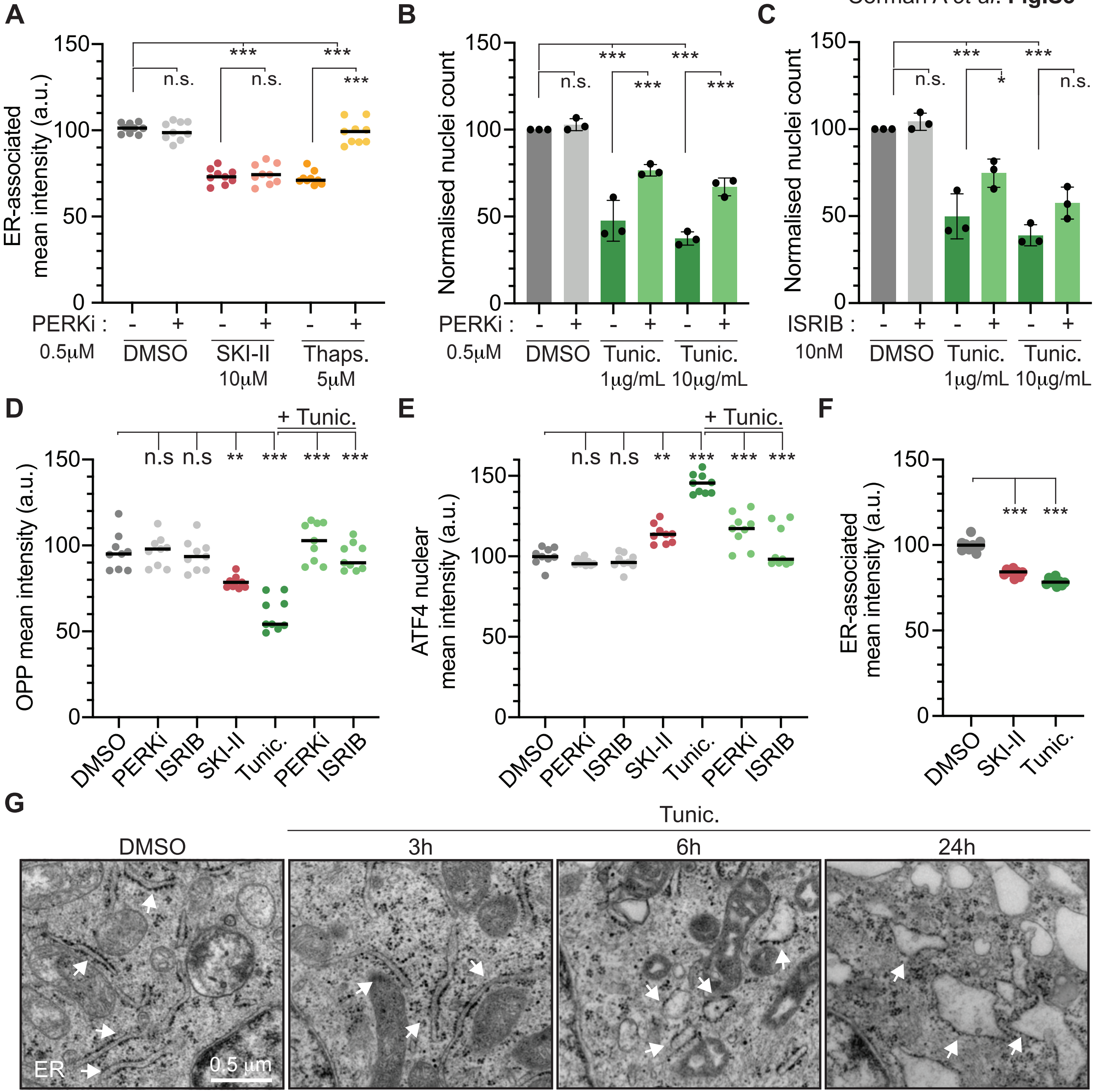

**Figure S3. Comparison of the effects of SKI-II to the ISR activating compounds Thapsigargin and Tunicamycin.** (**A**) HTM-dependent quantification of ER-associated mean intensity in U2OS cells exposed to PERKi (0.5 μM) for one hour and then to SKI-II (10 μM) or Thapsigargin (Thaps., 5 μM) for additional 3h. DMSO was used as a control. ***$p < 0.001$. (**B,C**) HTM-dependent quantification of nuclei counts (normalised to DMSO) of U2OS cells exposed to Tunic. (1 and 10 μg/mL) alone or Tunic. together with PERKi (0.5 μM) (**B**) or ISRIB (**C**) for 24h. ***$p < 0.001$. (**D**) HTM-mediated quantification of OPP signal in U2OS cells pre-exposed for 1h to PERKi (0.5 μM), ISRIB (50 nM) or DMSO, and then exposed to Tunicamycin (Tunic., 10 μg/mL) or DMSO for 3h. Treatment with SKI-II (10 μM) was included in the assay. ***$p < 0.001$. (**E**) HTM-quantification of ATF4 nuclear signal in U2OS cells treated as in (**C**). (**F**) HTM-mediated quantification of ER-associated mean intensity in U2OS cells exposed for 3h to Tunic- (10 μg/mL), SKI-II (10 μM) or DMSO. ***$p < 0.001$. (**G**) Representative TEM images of U2OS cells treated with Tunic. (10 μg/mL) for 3, 6 and 24h. Cells were exposed to DMSO as a control. White arrows point at the ER. Scale bars, 0.5 μm.
